# Mesoscopic cortical network reorganization during recovery of partial optic nerve injury in GCaMP6s mice

**DOI:** 10.1101/2020.06.28.173799

**Authors:** Marianne Groleau, Mojtaba Nazari-Ahangarkolaee, Matthieu P. Vanni, Jacqueline L. Higgins, Anne-Sophie Vézina Bédard, Bernhard A. Sabel, Majid H. Mohajerani, Elvire Vaucher

## Abstract

As the residual vision following a traumatic optic nerve injury can spontaneously recover over time, we explored the plasticity of cortical networks during the early post-optic nerve crush (ONC) phase. Using *in vivo* wide-field calcium imaging on awake Thy1-GCaMP6s mice, we characterized resting state and evoked cortical activity before, during, and 30 days after ONC. The recovery of monocular visual acuity and depth perception was evaluated at the same time points. Cortical responses to an LED flash decreased in the contralateral hemisphere in the primary visual cortex and in the secondary visual areas following the ONC, but was partially rescued between 3 and 5 days post-ONC, remaining stable thereafter. The connectivity between visual and non-visual regions was disorganized after the crush, as shown by a decorrelation, but correlated activity was restored 30 days after the injury. The number of surviving retinal ganglion cells dramatically dropped and remained low. At the behavioral level, the ONC resulted in visual acuity loss on the injured side and an increase in visual acuity with the non-injured eye. In conclusion, our results show a reorganization of connectivity between visual and associative cortical areas after an ONC, which is indicative of spontaneous cortical plasticity.

## Introduction

Recovery of vision after damage to the visual pathway in adults can happen both spontaneously and following a visual training, as long as there is still some residual vision^1–3^. After an optic nerve injury, for example, when retinal ganglion cells or their axons degenerate, some residual optic nerve fibers still connect to the superior colliculus and lateral geniculate nucleus of the thalamus^4–6^. The partial optic nerve crush (ONC) in adult rodents can be used as a model to simulate the visual deficits of diseases such as traumatic optic neuropathy, glaucoma, optic neuritis or immune-mediated CNS demyelination^7^, and its intensity determines the degree of vision loss. Spontaneous visual recovery occurs in this model with up to 20% gain of function, as shown in a contrast discrimination task or visually guided behavior in rats and mice^5,8–10^. The extent of residual vision depends on the number of surviving retinal cells and their axons, as well as cortical plasticity mechanisms, including functional connectivity reorganization^5,11^.

In fact, the dynamics of cortical network reorganization in central visual system structures are important mechanisms of visual recovery via both functional reconnection of lateral and recurrent circuits, and learning mechanisms, such as synaptic consolidation and synaptogenesis. Electrophysiological recording of individual neuronal activity, optical imaging of dendritic spines dynamics, or immunostaining methods, provided evidence for structural and functional changes in the lesion projection zone in the visual cortex, emerging a few hours/days after a retinal lesion or monocular enucleation^10,12–15^. However, it is still unclear how the visual pathways beyond the primary visual cortex (V1) respond to the loss of retinal input.

In the present study, we longitudinally monitored the residual cortical function and plasticity following an ONC over time in adult mice to characterize the spontaneous recovery of vision. To this end, we used wide-field calcium imaging on head-fixed awake mice expressing a calcium marker (Thy1-GCaMP6s) to assess how ONC affects the regional spread of sensory activation within both hemispheres. The response to a monocular flash illumination was measured before and at different time points after the ONC (1h, 1, 3, 5, 7, 14, 23, and 28 days post-ONC) in nine cortical structures: V1, secondary visual areas (A, PM, AM), the multimodal restrosplenial cortex (RS), and control areas not directly receiving retinal input, i.e. the anterior cingulate cortex (AC), primary motor cortex (M1) and somatosensory cortex (barrel cortex, BC, and Hindlimb sensory cortex, HL) (Fig. 1A). Patterns of spontaneous activity at rest were also assessed before and one month after ONC. Visual acuity and visual acuity-related depth perception were measured using the optomotor reflex response and visual cliff tests before and at the same time points after ONC. The extent of the retinal ganglion cell loss was quantified by immunohistochemistry. We observed a significant and widespread impairment of the response to a unilateral flash followed by a partial recovery of function in the visual cortical areas as early as the fifth day. The strongest effects were limited to the visual cortical areas (V1 and adjacent secondary areas) and RS cortex. The connectivity between visual and non-visual regions was however disorganized after the crush, as shown by decorrelation, but correlated activity was restored 30 days after the injury.

**Figure 1.**
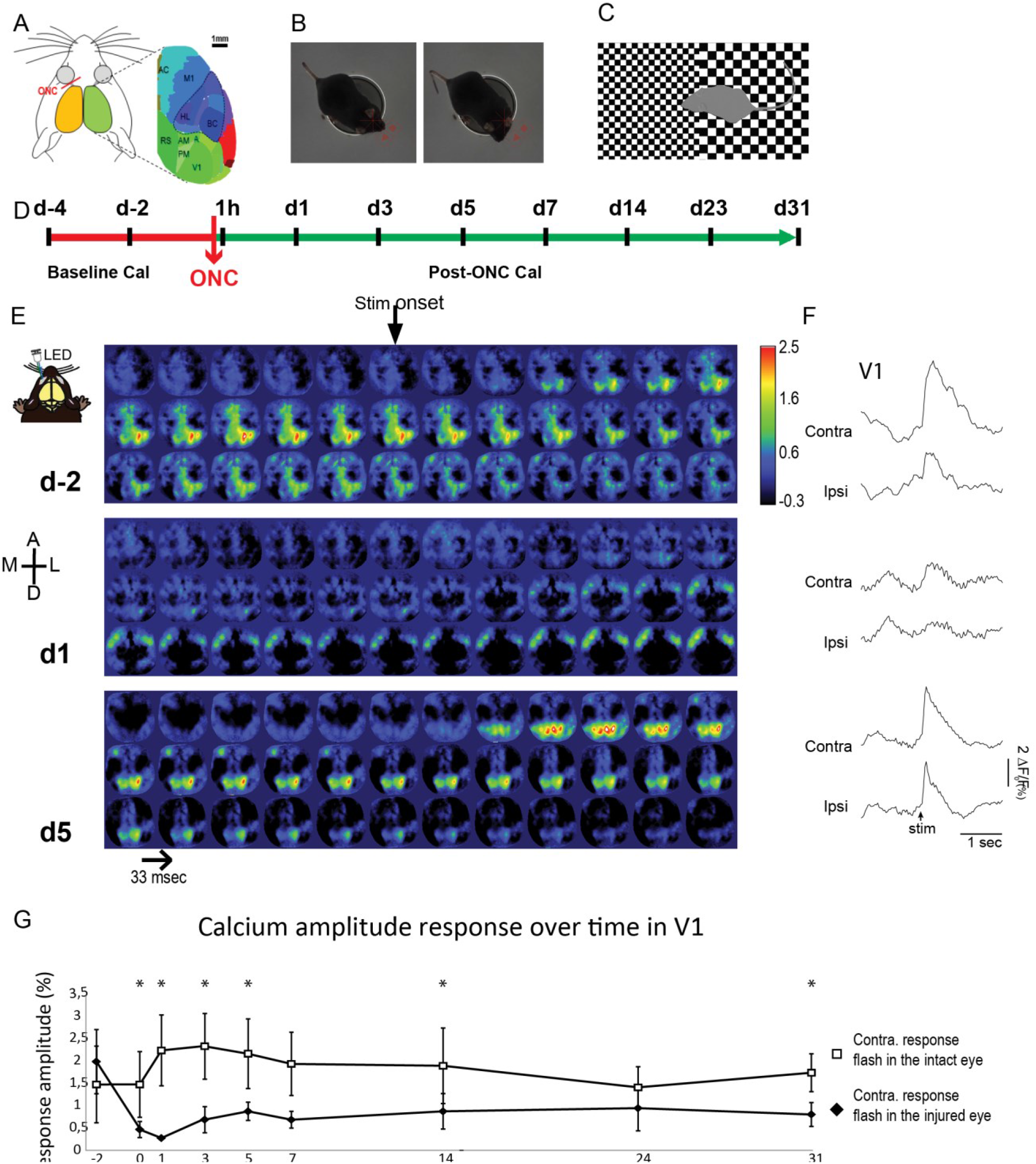
Mapping of a flash evoked response by wide-field calcium imaging before and after the optic nerve crush. Upper panel represents the t**imeline of the experiments**. **A**. Representation of the mouse head and brain showing the nine cortical regions of interest related to vision (V1, A, AM, PM, RS) or non-visual regions (M1, HL, BC) used to measure the neuronal activity by calcium imaging before and after partial optic nerve crush (ONC). **B**. Behavioral assessment of the tracking reflex in response to a drifting pattern as shown by two consecutive photographs of the mice. The arrow represents the drifting direction, the tracking reflex is shown by a movement of the head **C.** Behavioral assessment of depth avoidance with the visual cliff test in which the time spent by the mouse in the shallow vs deep end of an open field was recorded. **D.** After the implantation of an imaging chamber and habituation to the imaging setup, two baseline and eight post-ONC calcium imaging measurements were performed (after 1hour or at day 1, 3, 5, 7, 14, 23, 31) during the spontaneous recovery period. CaI, Calcium Imaging; V1, primary visual area; A, PM, AM, anterior, posterior medial, and anterior medial secondary visual areas; RS, restrosplenial cortex; AC, anterior cingulate cortex; M1, primary motor cortex; BC, barrel cortex; HL, Hindlimb sensory cortex. Lower panels represent the temporal decourt maps **E**. Maps at day −2 (d-2), day 1 (d1), and at day 5 (d5). The evoked response is contralateral to the LED flash. A round optical chamber was placed over both hemispheres over a denuded skull and a coverslip was added. Calcium imaging was performed through a CCD camera in head-fixed awake Thy1-GCAMP6s mice. **F** The panels to the right shows the amplitude vs time graphs in the left and right visual cortex (V1). **G.** Lower panel shows the response amplitude in the ipsilateral vs contralateral hemisphere of the mice during the spontaneous recovery period.

## RESULTS

Various parameters for the fluorescent calcium signal (ΔF/F_0_ × 100) were measured at different time points (pre-ONC, 1h, 1, 3, 5, 7, 14, 23, and 28 days post-ONC): the *amplitude*, the *latency*, and the *persistence* of cortical activation (area under the curve)^16,17^. These parameters are indicative of the strength, speed, and efficiency of the neural processing, respectively. For the sake of clarity, only the means and standard deviations of the amplitude and peak latency values significantly differing from pre-ONC are provided (Fig. 1, 2).

**Figure 2.**
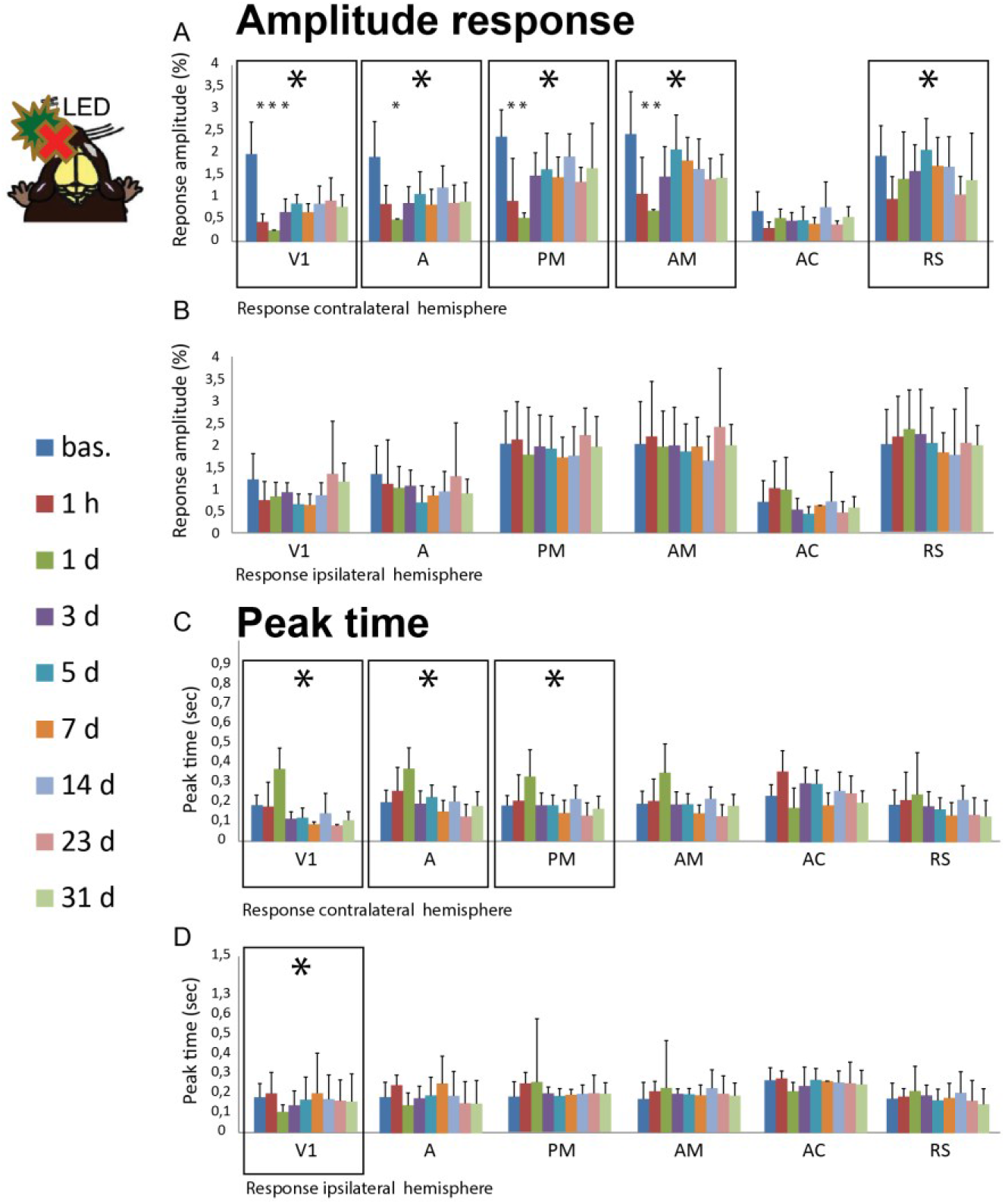
Cortical responses following optic nerve injury. Upper panel for amplitude and lower panel for peak response. **A.** Cortical peak response ΔF/F_0_ (%) in the contralateral hemisphere to the flash stimulation in the injured eye; **B.** Cortical peak response ΔF/F_0_ (%) in the ipsilateral hemisphere to the flash stimulation in the healthy eye; **C**. Peak latency in the contralateral hemisphere to the flash stimulation in the injured eye; **D.** Peak latency in the ipsilateral hemisphere to the flash stimulation in the healthy eye. V1, primary visual cortex; A, AM, PM, anterior, anteromedial, and posteromedial regions of the secondary visual cortex; AC, anterior cingulate cortex; RS, retrosplenial cortex.

### Pre-ONC responses to a flash

In naïve animals, the monocular flash LED stimulation (470 nm, 65 lm, 5 msec) induced a strong and transient increase in the calcium signal in the contralateral hemisphere to the stimulated eye (Fig. 1, 2). The strongest responses were seen in visual areas V1, A, PM, and AM. Similar to previous findings^16,17^, detectable but low responses were seen in other contralateral cortical areas (motor, sensorimotor, and cingulate cortices), as well as in the visual areas of the ipsilateral cortex, the latter probably being due to trans-callosal connections or retinal projections to the ipsilateral hemisphere.

### Partial cortical recovery of the amplitude of the response in the contralateral hemisphere following ONC

A Kruskal-Wallis test was conducted to examine the effect of an ONC on the amplitude of calcium responses, followed by a multiple comparisons test. The cortical response to a flash stimulation administered to the crushed eye was immediately decreased in the contralateral V1, A, AM, PM, and RS, cortical regions related to the visual pathway (n = 9, Fig. 3). In V1 (p = 0.000), the calcium response was lower at 1h (p = 0.000), 1 day (p = 0.000), and 3 days post-ONC (p = 0.034) compared to pre-ONC values (Fig. 2A). However, some cortical activity was recovered between 3 and 5 days post-ONC and remained stable thereafter until one month, when the experiment ended. The same tendency was observed in area A (p = 0.012), where a decrease was observed at 1 day (p = 0.011), in PM (p =0.000) at 1h (p = 0.012) and 1 day (p = 0.000), in AM (p = 0.001) at 1h (p = 0.026) and 1 day (p = 0.001) and in RS (p = 0.006). No change was observed in areas AC (p = 0.263), HL (p = 0.397), BC (p = 0.651), and M1 (p = 0.389) or in the ipsilateral hemisphere (Fig. 2B).

**Figure 3.**
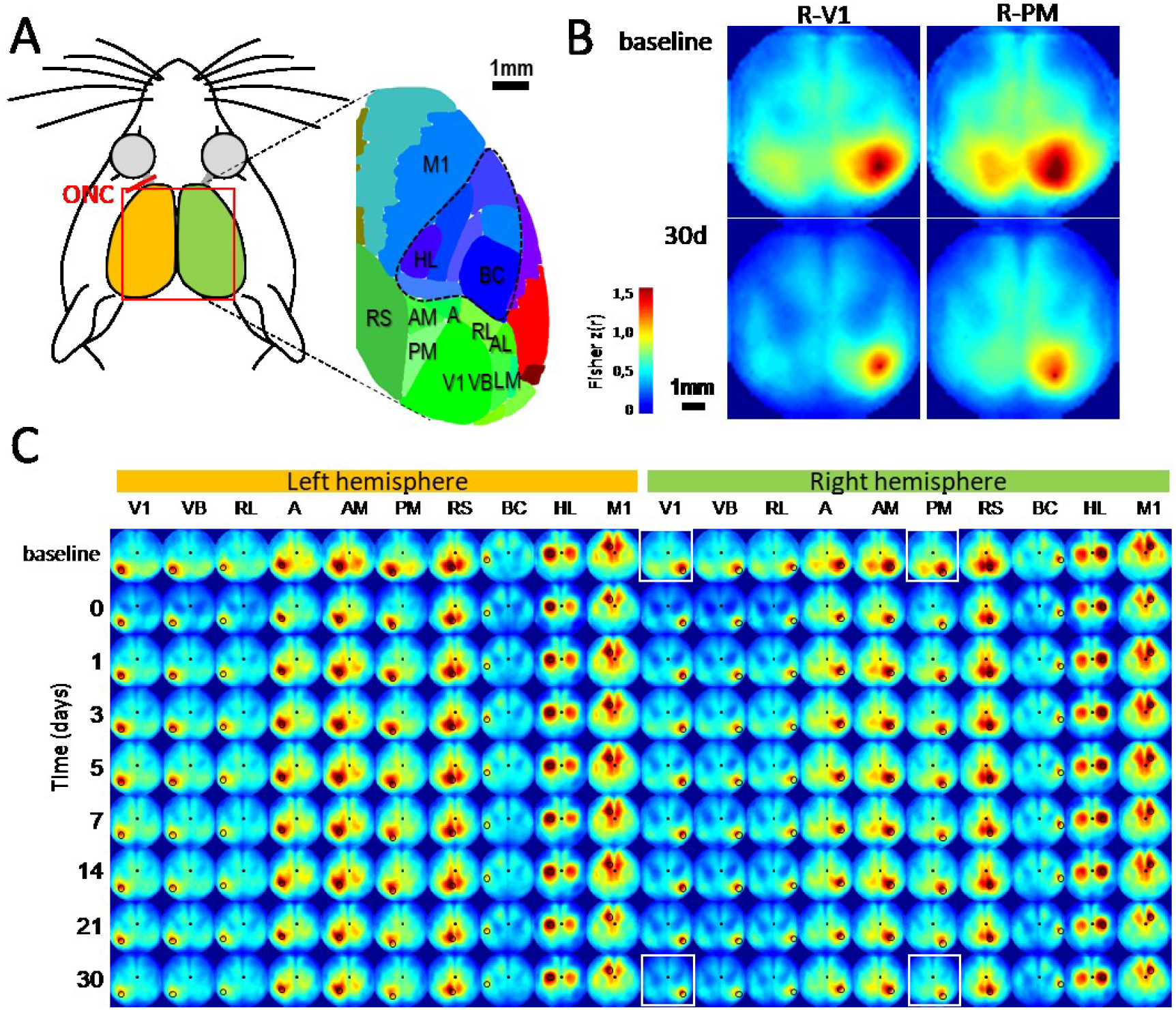
Seed Pixel Correlation Mapping. **A.** Schematic of the murine cortical regions imaged in the present study. **B.** Representative examples of Seed Pixel Correlation maps (averaged from n=9 mice and Fisher Z-transformed) for seed pixels localized in right areas V1 and PM before ONC (baseline) and 30 days after. **C.** Averaged Seed Pixel Correlation maps of 20 different locations (black circles) before and after the ONC (y-axis, 9 measures for 30 days). Correlation values were Fisher Z-transformed and displayed between z=0 and z=1.5 corresponding approximately to r=0.91.

The peak cortical response to the visual stimulation of the intact eye was not significantly affected by the crush of the opposite optic nerve in any regions, in either hemisphere (p > 0.05 for each region, compared to baseline, Suppl. Fig. 1).

### Rapid compensation in the efficiency of the response (peak latency) in both hemispheres

Further analysis response rapidity was conducted by comparing the latency of the maximal Ca^2+^ response to the onset of the flash using a Kruskal-Wallis test (Fig. 2C, D). The peak latency was delayed, subsequent to optic nerve injury, within V1 (p = 0.005) and secondary visual areas A (p = 0.004) and PM (p = 0.024) of the contralateral hemisphere during the first day after the ONC. However, 3 days after injury, the response in V1 was significantly more rapid over time and returned to pre-injury values in V2 of the ipsilateral hemisphere. No changes were seen in the upstream areas AM (p = 0.072), AC (p = 0.232), and RS (p = 0.490). In the ipsilateral V1 (p = 0.014), an increased latency of the visual stimulus response occured at day 1, returning to baseline values thereafter, given the shortest latency of the peak response after the flash. No changes were observed in ipsilateral areas A (p = 0.084), PM (p = 0.390), AM (p = 0.107), AC (p = 0.485), and RS (p = 0.283), or in any of the other non-visual cortical areas observed. The peak response time to the stimulation in the intact eye was not affected in either of the two hemispheres. The efficiency of cortical activation (peak duration) was not affected by the ONC at any time points.

### Comparison of the response in V1 of both hemispheres

Before the ONC, the calcium signal in response to a flash in the opposite eye was similar in the two hemispheres before the ONC. There was a significant difference in the activity of ipsilateral vs contralateral V1 following the visual flash stimulation in the intact (Fig. 1 D, white curve) or injured eye (Fig. 1 D, black curve) in the first 5 days (student t-test, p <0.05) following the ONC. After this early period, a partial recovery of the amplitude of the cortical response was observed, reducing the significance of the difference between responses in the cortical regions.

### Resting state spontaneous activity before and after the ONC

Spontaneous resting state activity was used to map the functional connectivity of the whole dorsal cortex (Fig. 3). As previously described^16–18^, during the baseline, seed pixel correlation maps showed a specific pattern of local and long-range connections between different visual areas and other visual regions (Fig. 3 B). Strong ipsilateral interconnections were also established between M1 and BC (reciprocally, Fig. 3 C). In addition, strong interhemispheric connections were also observed between bilateral RS, as well as M1 and HL homotopic areas.

One day after the ONC, a strong transient reduction of the overall connectivity was observed between all visual regions contralateral to the ONC, as well as some within the ipsilateral cortex (Fig. 4 A, B). A strong and long-term reduction of interhemispheric connections was observed between visual regions after 30 days, particularly between homotopic AM or PM areas (Fig. 4 C), but not in control regions such as homotopic HL, BC and M1. A strong decrease in local correlations (surrounding each seed) was also observed for the majority of visual regions in the contralateral cortex to the injured eye, but less in ipsilateral visual regions and other bilateral reference regions. This was particularly obvious between areas V1 and A, as well as between areas AM and PM. Additionally, we noted no increase of connectivity in any of the explored seeds.

**Figure 4.**
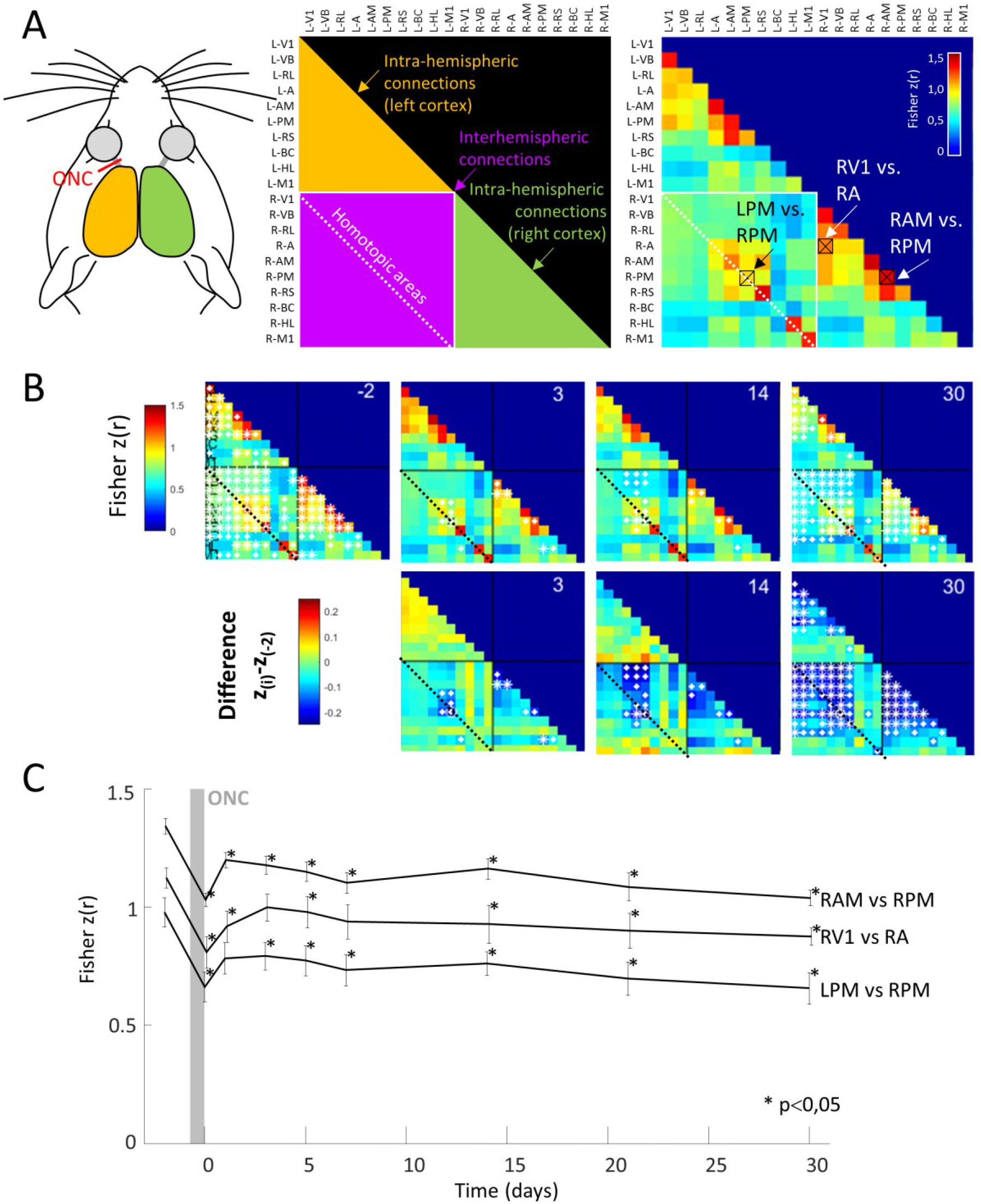
Functional connectivity matrix. **A.** *Left:* Schematic of connectivity information provided by the correlation matrix. L- and R-indicates left and right hemispheres respectively. Yellow and green zones: correlation values between regions of the same hemisphere (left and right respectively). Purple zone: correlation values between regions of both hemispheres. Dotted white line: correlation between homotopic regions from the two hemispheres. *Right:* Correlation matrix (averaged from n=9 mice and Fisher Z-transformed) during baseline with 3 representative connections used in panel C (black boxes). **B.** *First line:* Seed Pixel Correlation matrix averaged from n=9 mice for the 9 days before and after ONC. Size of white stars indicate the p value of the statistical comparison (small stars: p<0.05, medium: p<0.01, big: p<0.001). *Second line*: Differential Correlation Matrix (difference in correlation between each day and the baseline). **C.** Evolution of the correlation for 30 days (9 measures) in 3 representative connections (right areas AM and PM, right areas V1 and bilateral areas PM, as shown in panel A). (*) indicates Wilcoxon test, between baseline and current day and which provided p values under 0.05.

### The ONC altered visual function at the behavioral level

At day 1 post-unilateral ONC, a significant decrease in visual acuity in the lesioned eye (Wilcoxon, *p* = 0.012) was observed and remained stable up to one month following the injury. We found no recovery of visual acuity with the behavioural test over time following the ONC (Fig. 5A). However, an unexpected compensatory mechanism seems to occur in the intact eye, as evident by an increase in visual acuity over time after the ONC (Friedman *χ*^2^ = 20.454 *p* = 0.002) as of day 1 (Wilcoxon, *p* = 0.012) that remained up to 28 days (*p* < 0.05).

**Figure 5.**
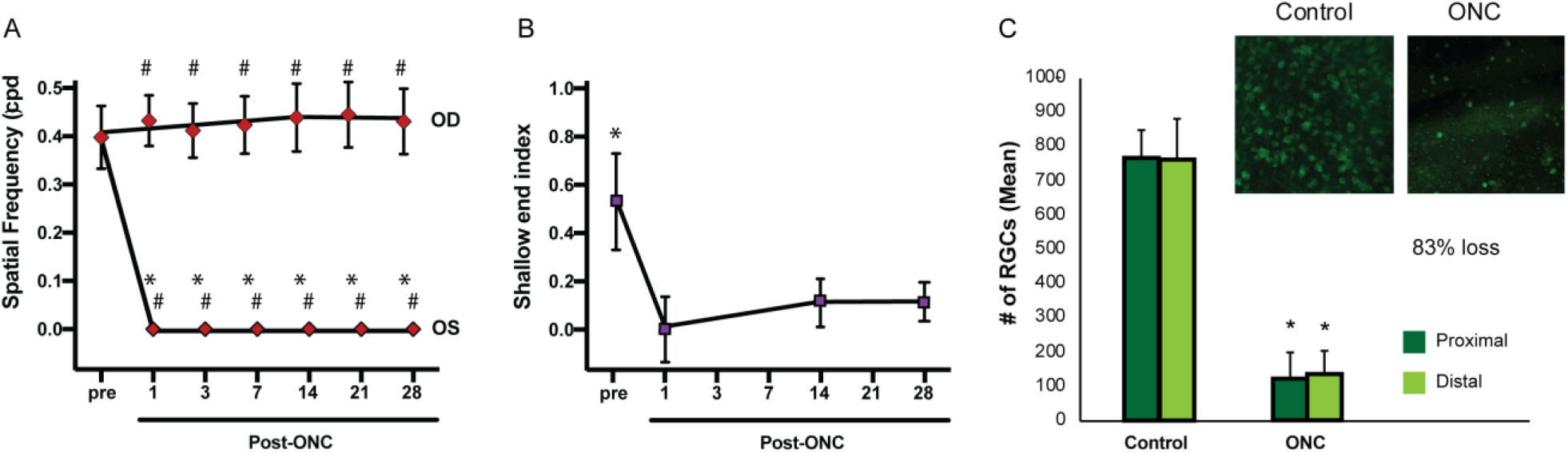
Effect of ONC on visual acuity and retinal cell loss. **A** Following the optic nerve injury (day 1–28), there was a significant decrease in visual acuity of the injured eye and a slight increase of visual acuity of the intact eye when compared to the baseline values. **B:** Following the optic nerve injury (day 1–28), there was a significant decrease in depth perception, which induced a decrease in time spent in the shallow end with no significant recovery. **C** The ONC induced a severe loss of retinal ganglion cells (83%) both in central (proximal, dark green) and peripheral (distal, ligth green) regions of the retina. Insert: microphotographs of flat mount retinas stained with RBPMS. ONC: optic nerve crush.

A complementary behavioral test evaluating visual depth discrimination, a cortical function, was also performed (using the bilateral ONC). Prior to the ONC, the mice had a significant preference for the shallow end during the visual cliff test (Wilcoxon, *p* = 0.018), demonstrating hesitancy and avoidance behaviors toward the deep end (Fig. 5B). The preference for the “shallow end” did not return in the following 28 days (Friedman, *χ*^2^ = 1.143, *p* = 0.565)

### Survival of retinal ganglion cells after ONC

For the unilateral ONC, RGC loss was significant in the damaged eye compared to the non-injured eye in both proximal and distal regions (Mann-Whitney, *p* = 0.000). One month after the ONC, a survival rate of approximately 10% of RGCs was found in the injured eye (Fig. 5C). This indicates that, despite the low survival of RGCs at one month following the lesion, the survival rate is higher than what is found in an axotomized retina. In the bilateral ONC group, the RGC survival was about 17%, with no significant difference between proximal and distal distributions (Mann-Whitney, *p* = 0.008).

## DISCUSSION

Our study was designed to gain a better understanding of how the cortex reorganizes itself over time, following a partial ONC using mesoscopic mapping, and how this relates to the recovery of vision. A significant and widespread impairment of the response to a unilateral flash was observed in all cortical areas, which was followed by a partial restoration of the cortical visual activity as soon as the fifth day and some acuity change of the opposite eye. The effects were limited to the visual cortical areas (V1 and adjacent secondary areas) and RS. Moreover, the response latency of V1 was enhanced post-injury compared to pre-ONC, suggesting a reduction of input processing complexity in V1. The connectivity between visual and non-visual regions was disorganized after the crush, as shown by a decorrelation, but correlated activity was restored 30 days after the injury.

### The ONC induces a visual impairment followed by a gradual recovery of cortical activity

A partial recovery of the cortical activity was observed post-ONC, subsequent to an initial drop of the cortical functioning, in terms of amplitude, reactivity, and co-activation. The visual cortex is at first inactivated by the lack of retinal input and related impairment^19^, similar to the decrease in brain function following optic neuritis, which is proportional to the extent of the damage of the optic nerve^20–22^. These results are in line with recent findings showing cortical plasticity in V1 after an ONC in rats or mice^10,23–25^ followed by a recovery of cortical activity. This recovery is in agreement with previous autoradiographic studies after an ONC in rats^26^. Other studies were unable to observe an improvement in cortical activity following an ONC using visual evoked potential (VEP) recordings^9,27,28^ in anesthetized rats. The calcium signal of GCAMP6s mice mainly arises from the Thy-1 long-projecting excitatory neurons and is not detected in GABAergic cells^29^. Due to the density of the cortical tissue, the fluorescent signal from the superficial layers would have a stronger influence on the acquired signal compared to the signal from deeper layers, which would be more diffuse^30^. This signal might be stronger than VEP recordings from layer 4 in terms of cortical response following an ONC. A layer-specific response to sensory stimulation has been confirmed by 3-photon calcium imaging^31^. This suggests that the hyperactivity in layer 4 due to reduced visual inputs that impair VEPs may be compensated in layers 2/3 by corticocortical connections and recurrent circuits. Moreover, synaptic plasticity or dendritic spine density has also been shown to be involved in the recovery of activity in V1^15,32^. However, it is not known whether this recovered activity is enough to trigger spiking activity, although the response amplitude in other cortical areas was not affected by the visual differentiation. It is interesting to note that ocular dominance might also be affected by the crush, as the response in the non-affected cortex increased compared to the opposite cortex, which agrees with a recent study showing change in the ocular dominance index in the binocular cortex^10^.

Recovery of function has also been demonstrated by optical imaging after a retinal lesion^15^. It was found that a focal retinal lesion produces an anatomically defined lesion projection zone, which is rapidly and functionally restored due to long-range horizontal fibers projecting to the borders and center of the lesion projection zone^14,33–36^. In contrast, our ONC model causes a degeneration of ganglion cells, which is randomly distributed with approximately 10% retinal ganglion cell survival, i.e. it is a reduction of visual input that is randomly distributed throughout the entire extent of V1. Spontaneous recovery of the calcium signal in V1 most likely results from a combination of residual inputs from the surviving cells and cortical plasticity arising from strengthened cortico-cortical or local lateral connections^33,35,36^, including cross-modal innervation^37^. Cortical reorganization after optic nerve or retinal lesions might be altered at the level of long-range lateral connections, as well as the global functional connectivity network^34,38,39^. This would be in line with human studies where patients with optic nerve injuries had not only functionally disturbed connectivity networks^40^, but also subtle deficits in the presumably “intact” areas of the visual field^41^.

### Plasticity of the functional connections after ONC

To determine whether long-range corticocortical or local connections were involved in the recovery of function, or if other cortical structures would compensate for the reduced functioning in V1, we used seed pixel correlation mapping based on resting state spontaneous activity^21,43^. We observed a strong transient reduction of the overall connectivity within the bilateral visual cortex shorty after the ONC. These acute side effects of the ONC surgery should have a minimal effect on transient loss of connection at Day 0 because some were preserved within the somatosensory and motor cortices. This short-term effect was followed by a clear long-term reduction of the connections between homotopic extra striate areas, as well as between area V1 and some extra striate areas of contralateral hemisphere to the ONC. Long-term changes between extra striate areas were also observed. Surprisingly, no increase in connections was observed among other regions to compensate for the loss of these connections. Overall, these results showed that most of visual areas decrease their long-range connections and establish more local computations. This local cortical effect and local reorganization of sensory deprivation is distinctly in contrast to other losses of function such as a transient focal stroke that globally depresses cortical activity, followed by a circuit reorganization and functional remapping within the entire network^42^. Hind limb ligation also resulted in large-scale reductions in functional connectivity that were not restricted to the hind limb primary somatosensory region^43^. Our results suggest that the V1 is the main permissive gating area for visual input processing.

### The ONC induces visual behavioral alterations

Visual function decreased substantially following the ONC, as measured by the optomotor reflex and visual cliff tests. The ability to track moving sinusoidal gratings was no longer observed via the optomotor reflex test after ONC. Similarly, our mice did not appear to detect the cliff as indicated by the loss of avoidance behaviors during the visual cliff test. Previous rat studies showed a disconnection between behavior and anatomical or electrophysiological changes following damage to the optic nerve^9^. In these studies, it was shown that even with only 10% of functional retinal ganglion cells, the animals were able to perform close to normal in behavioral visual tasks^26^. In our study, it took approximately 5 days to observe a partial recovery of cortical activity following the ONC using the calcium imaging technique, but no recovery of visual perception was noted in the lesioned side. This might be due to the intensity of the crush, which was stronger than what was used in the Schmitt and Sautter studies, as their lower ONC intensity allowed for greater recovery. Similarly, in another study, a residual cell number of 30% supported the recovery of visual acuity and discrimination performance three weeks after the ONC (data not shown). It is also possible that our tests measure finer visual capacities compared to light discrimination tests used in the previous studies mentioned above. The discrepancy between the recovery of visual acuity in the non-deprived eye and the absence of apparent cortical activity on the ipsilateral cortex is also puzzling, suggesting that the recovery of visual acuity measured by the optomotor reflex mostly relies on subcortical circuits. It has already been proposed that this test depends on the integrity of the superior colliculus, which is involved in the control of saccades and head movements^5^.

### Limitations

It is possible that a limitation of our model was the use a high intensity ONC, whereas a lower intensity would promote better perceptual recovery as well as increased amplitude of the response. This would fit with a computational study demonstrating a correlation between the extent of recovery and the sensory lesion size^44^. Furthermore, the decrease of brain function in humans following optic neuritis has been correlated with the extent of the damage occurring in the optic nerve^20–22^. Accordingly, a low intensity ONC in mice allows for partial recovery of behavioral discrimination (data not shown). However, the partial recovery of neuronal activity with a strong intensity crush is very interesting.

## Conclusion

In conclusion, our results show a reorganization of the local connectivity between visual cortical areas following a traumatic optic nerve injury. This is indicative of visual cortical plasticity, though it does not extend to higher order cortical areas during the short recovery period.

## Methods

### Animal preparation

All procedures were carried out in accordance with the guidelines of the Canadian Council for the Protection of Animals and were accepted by Animal Welfare Committee of the University of Lethbridge and by the Ethics Committee of the Université de Montréal (protocol CDEA 17-010). A total of 9 C57BL/6 J-Tg(Thy1-GCaMP6s)GP4.3Dkim/J adult mice (IMSR Cat# JAX:024275, RRID:IMSR_JAX:024275; 3 females, 6 males; Jackson Laboratory, Bar Harbor, ME, USA) were used for the imaging in this study. Moreover, 9 C57BL/6NCRL males (IMSR Cat# CRL:027, RRID:IMSR_CRL:027) from Charles River Laboratories (RRID:SCR_003792) were tested for their optomotor reflex after a unilateral ONC, and 7 C57BL/6NCRL males were used for the optomotor reflex and visual cliff testing for the bilateral ONC. All animals were maintained in a 12 h light/dark normal daylight cycle with *ad libitum* access to food and water.

### Imaging chamber implantation

Mice were anesthetized with isoflurane (3.5% induction, 1.5% maintain). After a subcutaneous (s.c.) injection of 500 μL of glucose buffers, the scalp was disinfected with iodine, and then lidocaine was injected subcutaneously. Core body temperature was maintained at 37°C using a feedback-controlled heating pad. The skin covering the skull was entirely removed and the skull was allowed to dry. A cover slip was placed over both hemispheres and was secured with transparent dental cement (C&B MetaBond, Parkell, Edgewood, NY, USA). A custom-made round imaging chamber with two arms (titanium; 10 mm diameter) was also fixed to the skull.

### Optic Nerve Crush (ONC) surgery

A left partial optic nerve crush^45–47^ was performed under isoflurane anesthesia for the calcium imaging. For complementary behavioral experiments with the visual cliff, an ONC was performed on both eyes. Briefly, the optic nerve was exposed from the lateral side of the eye by an opening in the conjunctiva and then crushed with calibrated self-clamping forceps (Martin Instruments, Tuttlingen, Germany) at 1-2 mm from the eye for 3 sec. Retinal blood supply and dura were left intact. An ophthalmic antibiotic (Ciloxan^®^, Alcon Canada inc., Missisauga) was topically applied to the eye after the surgery.

### Calcium imaging

The neuronal activity was measured by calcium imaging according to two paradigms: during the resting state and during visual stimulation with a flash at different time points before and after ONC.

#### Apparatus set-up

The awake mice were restrained at the head, with the chamber arms fixed to two laterally placed clamps. They were positioned on a stage with adjusted height to give the best comfort to the animal. A 470 nm visual stimulation LED fixed on a 20 mm Star Base (Rebel LED, 65 lm, 700 mA, Luxeon) was placed at 5 cm from one eye. The mice were gradually habituated to the apparatus and the visual stimulation of 15 min/day for one week, so that they were not stressed and relatively quiescent during imaging. During head fixation, animals were placed in a plastic tube to limit motion and guide relaxation.

#### Visual stimulation

Visual stimulation consisted of a flash of 5 msec (65 lm), delivered 2 sec after the onset of imaging. This sequence was completed 30 times (total duration of 6 sec) at 10 seconds apart from each other in one eye, repeated thereafter in the other eye. Evoked responses were recorded at 30 frames per second for the duration of 6 sec per trial.

#### Image acquisition

The cortex was illuminated with a blue LED (Luxeon K2, 473 nm center) and excitation filter of 470 ± 20 nm. Emitted fluorescence was filtered using a 527 to 554 nm bandpass optical filter (Semrock). A CCD camera (1M60 Pantera, Dalsa) was controlled by an EPIX E8 frame grabber with XCAP 3.8 imaging software (EPIX, Inc.) to record optical images. Images were collected with a macroscope composed of front-to-front video lenses (8.6×8.6 mm field of view). Frames of 128×128 pixels were acquired at a rate of 30 Hz, giving a spatial resolution of 67 μm/pixel. To capture the fluorescence originating from the cortex, the focal plane was moved down ~1 mm from the skull surface. In the case of the resting state spontaneous activity data, the recordings were performed in absence of any visual stimulation for a 10 min period.

#### Data analysis

Calcium response to visual stimulation (flash) was calculated as the normalized difference to the baseline recorded 2 sec before stimulation (ΔF/F_0_ × 100) and averaged across 30 trials using ImageJ (ImageJ, RRID:SCR_003070; National Institutes of Health). The map was smoothed using a Gaussian spatial filter with a SD of 1 pixel. To characterize the changes in the visual responses after the ONC in different cortical areas, we divided both cortical hemispheres into 9 distinct 0.11 mm^2^ regions of interest (ROI) (Fig. 1). All cortical regions were defined based on the stereotaxic coordinates relative to Bregma, using the Franklin & Paxinos mouse brain atlas (<u>Mouse Brain Atlases</u>, RRID:SCR_007127) and cortical map from Allen Brain Institute (<u>Allen Reference Atlas – Mouse Brain</u>, RRID:SCR_013286). Amplitude of response was calculated for each ROI as the calcium signal peak value after visual stimulation. Peak time was calculated as the latency between the calcium signal peak and the onset of visual stimulation. Response amplitude and latency were calculated using a custom script in MATLAB (<u>MATLAB</u>, RRID:SCR_001622, Mathworks).

Resting state spontaneous activity was band pass filtered between 0.3 to 3Hz and Pearson correlations were used between the activity of 20 seeds and other pixels of dorsal cortex for 10 days. To create seed pixel correlation maps, the cross correlation coefficient r values between the temporal profiles of one selected pixel and all the others of the ROI were calculated^17,48^. Seed pixel correlation maps were spatially registered in reference to Bregma, which was averaged for all mice. Bilateral seeds were V1 & VB: monocular and binocular zones of the primary visual cortex; RL, A, AM, RL: medial extra striate areas RS: retrosplenial cortex; BC: barrel cortex; HL: somatosensory cortex hindlimb and M1: motor. Peasons r-values were transformed via Fisher Z in order to perform statistical comparisons between connections before (average between two baseline times) and after the ONC (restricted to correlation r-values higher than 0.5) using nonparametric Wilcoxon paired tests (signed rank), as the Kolmogorov-Smirnov analysis did not confirm normal distribution.

### Behavioral assessment of visual acuity

Two behavioral tasks were used to assess visual processing^6^ (i.e., retina-superior colliculus pathway, lateral geniculate-V1 pathway, accessory optic areas, higher cognitive areas, etc.). The rate of vision recovery and how it relates to metabolic changes at various cortical levels of the visual system was of particular interest.

#### <u>Visual acuity related to the retina-superior colliculus pathway</u> (Optomotor reflex)

Mice were individually placed in a virtual cylinder with sinusoidal drifting gratings (12.0 d/s) (**OptoMotry™ apparatus**, Cerebral Mechanics Inc)^49^ and the tracking reflex in response to the drifting pattern was monitored. Briefly, the apparatus consisted of four computer screens, forming a 360-degree visual stimulation. A camera in the apparatus relayed a live transmission to the OptoMotry™ HD 2.0.0 software, which allowed for the manipulation of stimuli and observation of the animal’s behavior. First, the mice were habituated to the apparatus over the course of three days (5, 10, 15 minutes), alternating gray screen and moving gratings. The apparatus was cleaned with water following each test. Once the habituation was complete, the mice were tested prior to the ONC and 1, 3, 7, 14, 21, and 28 days thereafter. During testing, each mouse was placed in the apparatus and presented with a 0.050 cpd sinusoidal grating for 3 sec. We observed whether the mice followed the movement of the grating with their head, which was the criterion for detected spatial frequency. We presented full contrast spatial frequencies moving in both clockwise and counter clockwise directions at a speed of 12.0 d/s. Each eye was tested independently according to the direction of the grating movement. Spatial frequencies were incrementally increased (from 0.05 to 0.7 cpd) in an adaptive staircase procedure until no tracking activity was observed, which determined visual acuity thresholds.

#### Visual cliff

Depth avoidance was evaluated in an open field (transparent box) to assess depth perception based on visual discrimination cue. The method was adapted from that described by Lima et al.^5^. Specifically, a plexiglass box (40 cm L x 10 cm W) was positioned on a table so that half of the box was positioned directly over a 60 cm-long, 2 cm x 2 cm checkerboard pattern, while the other half was suspended 70 cm above an identical checkerboard. For each test, the mouse was placed in the back of the “shallow end” and movements were recorded for two minutes. Since the floor was level, the mice had to rely on visual cues to distinguish the “shallow end” from the “deep end” (as they tend to naturally avoid cliffs). The total time in the “shallow end” was noted, and the box was cleaned with water between each assessment. This cliff test was performed once prior to the ONC, and 1, 14, and 28 days thereafter.

### Estimation of RGC survival after ONC

Whole-mounted retinas were prepared for immunostaining. At the end of the experiment, mice were perfused with 4% paraformaldehyde in 0.1M phosphate buffer sodium at room temperature under deep anesthesia (pentobarbital, 52 mg/kg, i.p.). Two small holes were made in the cornea and the eyes were post-fixated in 4% formaldehyde overnight. The cornea and lens were removed, and the retina was dissected ex vivo in four quadrants. Sections were pre-incubated overnight at 4° Celsius in phosphates buffer with 0.5% triton (PBS, 0.1 M, pH 7.4) containing 10% goat serum and 1% BSA. The tissue was incubated for 48h at 4° Celsius with anti-RBPMS primary antibodies (1:500, Phosphosolutions; Cat# 1832-RBPMS, RRID:AB_2492226) in PBS-triton-0.5 % with 3% goat serum and 1% bovine serum albumin. This was followed by a 48h incubation in goat antirabbit IgG H&L (Alexa Fluor 555) antibody (1:500, Abcam, Abcam Cat# ab150078, RRID:AB_2722519) and then placed for 1 h in a Hoescht:PBS (1:10 000) solution. Quantitative examination of RBPMS immunoreactivity was used for ganglion cell counting in 1 mm^2^ squares at four proximal and four distal positions to the optic nerve region.

### Statistics

Statistical analysis of the calcium signal and latency was performed using non-parametric Kruskal-Wallis tests with a significance level of p < 0.05. Pairwise comparisons were performed between the amplitude and peak time of the values of the different days and the baseline values.

To assess vision loss as quantified with the optomotor reflex test (both monocular and binocular), a related-sample Wilcoxon signed-rank test was used with a significance level of p < 0.05 to compare the pre and post ONC day 1 data. Then, a non-parametric Friedman test served to evaluate if there was significant recovery (p < 0.05) after the ONC. This test, if significant, was followed by a Wilcoxon signed-rank test as post-hoc with a significance level of p < 0.05 to determine if there was a significant difference between specific days. The visual cliff test was analyzed using the related samples non-parametric Wilcoxon signed-rank test to compare the time spent in the “shallow end” versus time spent in the “deep end”.

The independent samples, non-parametric Mann-Whitney U test was used between the normal retina and the lesioned retina for the proximal and distal cell count with a significance level of p < 0.05.

## Data availability

The datasets generated during and/or analyzed during the current study are available from the corresponding author on a reasonable request.

## Acknowledgments

We are grateful to the Canadian Institute of Health Research (Grant No. MOP-111003 to EV), the Natural Sciences and Engineering Research Council of Canada (Grant No. 238835-2011 to EV), and FRQS Vision Health Research Network (RNI grant to EV, BAS, MV and MHM).

## Author contributions statement

MG, MN, JLH, ASVB, MV: contributed to the acquisition, analysis, and interpretation of the data in this study. MG, EV, MV, BAS and MHM: contributed to design of this study and interpretation of the data; MG, JLH: drafted the first version of the manuscript, EV, MV: wrote the last version of the manuscript. All authors have approved the final version of the manuscript; agree to be accountable for all aspects of the work in ensuring that questions related to the accuracy or integrity of any part of the work are appropriately investigated and resolved; designated as authors qualify for authorship and who qualify for authorship are listed.

---

**Suppl. Figure 1.**
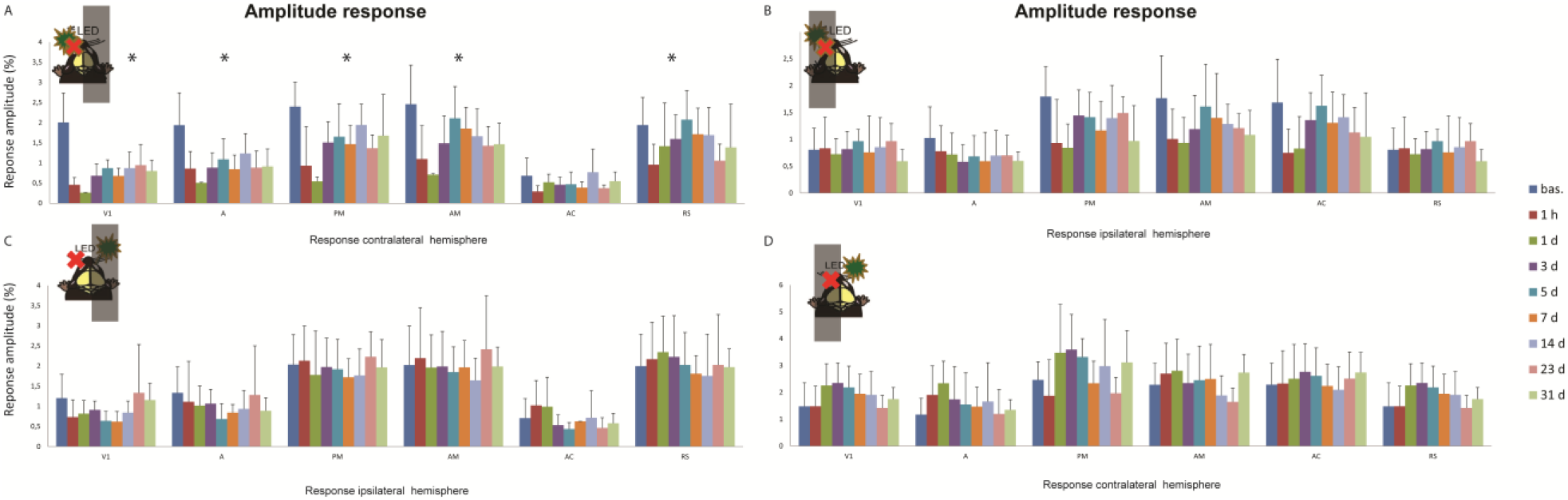
Amplitude responses in both hemisphere upon stimulation of the left or right eye following optic nerve injury. Upper panel for Cortical peak response ΔF/F_0_ (%) upon stimulation of the left eye and lower panel amplitude response upon stimulation of the left eye. **A.** Cortical peak response ΔF/F_0_ (%) in the contralateral hemisphere to the flash stimulation in the injured eye; **B.** Cortical peak response ΔF/F_0_ (%) in the ipsilateral hemisphere to the flash stimulation in the injured eye **C**. Cortical peak response ΔF/F_0_ (%) in the ipsilateral hemisphere to the flash stimulation in the healthy eye; **D.** Cortical peak response ΔF/F_0_ (%) in the contralateral hemisphere to the flash stimulation in the healthy eye. V1, primary visual cortex; A, AM, PM, anterior, anteromedial, and posteromedial regions of the secondary visual cortex; AC, anterior cingulate cortex; RS, retrosplenial cortex.

